# The qBED track: a novel genome browser visualization for point processes

**DOI:** 10.1101/2020.04.27.060061

**Authors:** Arnav Moudgil, Daofeng Li, Silas Hsu, Deepak Purushotham, Ting Wang, Robi D. Mitra

## Abstract

**Summary:** Transposon calling cards is a genomic assay for identifying transcription factor binding sites in both bulk and single cell experiments. Here we describe the qBED format, an open, text-based standard for encoding and analyzing calling card data. In parallel, we introduce the qBED track on the WashU Epigenome Browser, a novel visualization that enables researchers to inspect calling card data in their genomic context. Finally, through examples, we demonstrate that qBED files can be used to visualize non-calling card datasets, such as CADD scores and GWAS/eQTL hits, and may have broad utility to the genomics community.

**Availability and Implementation:** The qBED track is available on the WashU Epigenome Browser (http://epigenomegateway.wustl.edu/browser), beginning with version 46. Source code for the WashU Epigenome Browser with qBED support is available on GitHub (http://github.com/arnavm/eg-react and http://github.com/lidaof/eg-react). We have also released a tutorial on how to upload qBED data to the browser (dx.doi.org/10.17504/protocols.io.bca8ishw).

## Introduction

Advances in genomic technologies often lead to new data formats and new platforms to visualize those data. The Human Genome Project originated the popular Browser Extensible Data (BED) standard for describing genomic intervals (Kent et al., 2002). Routine next-generation sequencing projects, such as whole-genome sequencing and RNA-seq, use the SAM format to store and visualize data (Li et al., 2009). Epigenetic modifications detected by bisulfite sequencing can be visualized using methylC tracks (Zhou et al., 2014). The bedGraph and wiggle (.wig/.bigWig) formats have emerged as flexible standards for encoding pseudo-continuous integer- and real-valued signals across the genome, such as from normalized ChIP- or ATAC-seq assays (Huy Hoang and Sung, 2014; Kent et al., 2010; Rosenbloom et al., 2010). Finally, the .hic and .cool formats (Abdennur and Mirny, 2019; Durand et al., 2016) encapsulate intra-chromsomal contact frequencies and have contributed to our understanding of chromatin organization.

Over the past several years, we have introduced, and developed, transposon calling cards to identify genome-wide transcription factor (TF) binding sites (TFBS) (Wang et al., 2007, 2008, 2011, 2012). This approach uses a TF of interest fused to a transposase. The fusion construct deposits transposons into the genome near TFBS, which can be recovered from either DNA or RNA libraries. Significantly enriched clusters of transposons indicate putative TFBS. Instead of plotting read coverage, as would be done in more traditional TF studies like ChIP-seq, we plot each insertion as a discrete point along the (genomic) *x*-axis and the number of reads supporting that particular insertion on the *y-*axis. The result resembles a scatterplot in which an increased density of insertions is typically observed near TFBS.

Historically, raw insertion data were visualized using GNASHY, an in-house file format and genome browser custom built for calling card data. While useful, the GNASHY browser suffered from two major limitations: first, it was restricted to visualizing one track-and therefore, one sample or experiment-at a time; and second, it did not support conventional genomic formats like bedGraph or bigWig. Thus, any comparative analysis of calling card data with, say, ChIP-seq or ATAC-seq relied on manually aligning images from different browsers (Wang et al., 2012).

Calling card technology is currently undergoing a renaissance. We have recently used calling cards to study TF binding in bulk populations of cells *in vivo* (Cammack et al., 2020), and we have also combined calling cards with single cell RNA-seq to simultaneously profile cell identity in complex organs and heterogenous disease states (Moudgil et al., 2019). Calling cards has also been used to dissect TF binding in both steady state and dynamic contexts (Mayhew and Mitra, 2014; Shively et al., 2019). As the scope and application of the calling card technique grows, we anticipate greater interest and increasingly complex visualization demands. Here, to better support existing and future users, we describe the qBED format, a new text-based genomic data format for storing calling card data. We also describe the qBED track, an interface for visualizing calling card data on the WashU Epigenome Browser. Finally, we present examples of non-calling card genomic data visualized using the qBED standard to demonstrate the format’s flexibility.

## Implementation

We christened our format *q*BED because it stores multidimensional, *quantitative* information about *quantized* events, such as calling card transpositions. Formally, qBED follows a BED3+3 standard (Figure 1A). For calling card data, the first three columns denote the chromosome, start, and end coordinates of the transposon insertion. The width of the interval depends on the transposase used: mammalian calling cards, which employs the *piggyBac* transposase, uses a four base-pair width for the insertion coordinate as *piggyBac* overwhelmingly inserts into TTTA tetramers (Wang et al., 2012); whereas yeast calling cards uses single base-pair intervals as these assays use the motif-agnostic Ty5 retrotransposon (Wang et al., 2007). qBED files inherit the BED format’s 0-based, half-open intervals and are compatible with programs like bedtools (Quinlan and Hall, 2010) and bedops (Neph et al., 2012) for intersection analysis. The fourth column encodes a numerical value-in (in this case, the number of reads supporting each insertion–and is the last column required in qBED files. The fifth and sixth columns are optional, but recommended, as the former denotes the strand (+/-, or . if unspecified) that was targeted, while the latter encodes an annotation string. For calling card experiments, this is where sample-specific barcodes are registered (Figure 1A). Like BED files, qBED files can be compressed and indexed with bgzip and tabix, respectively (Li et al., 2009).

**Figure 1:**
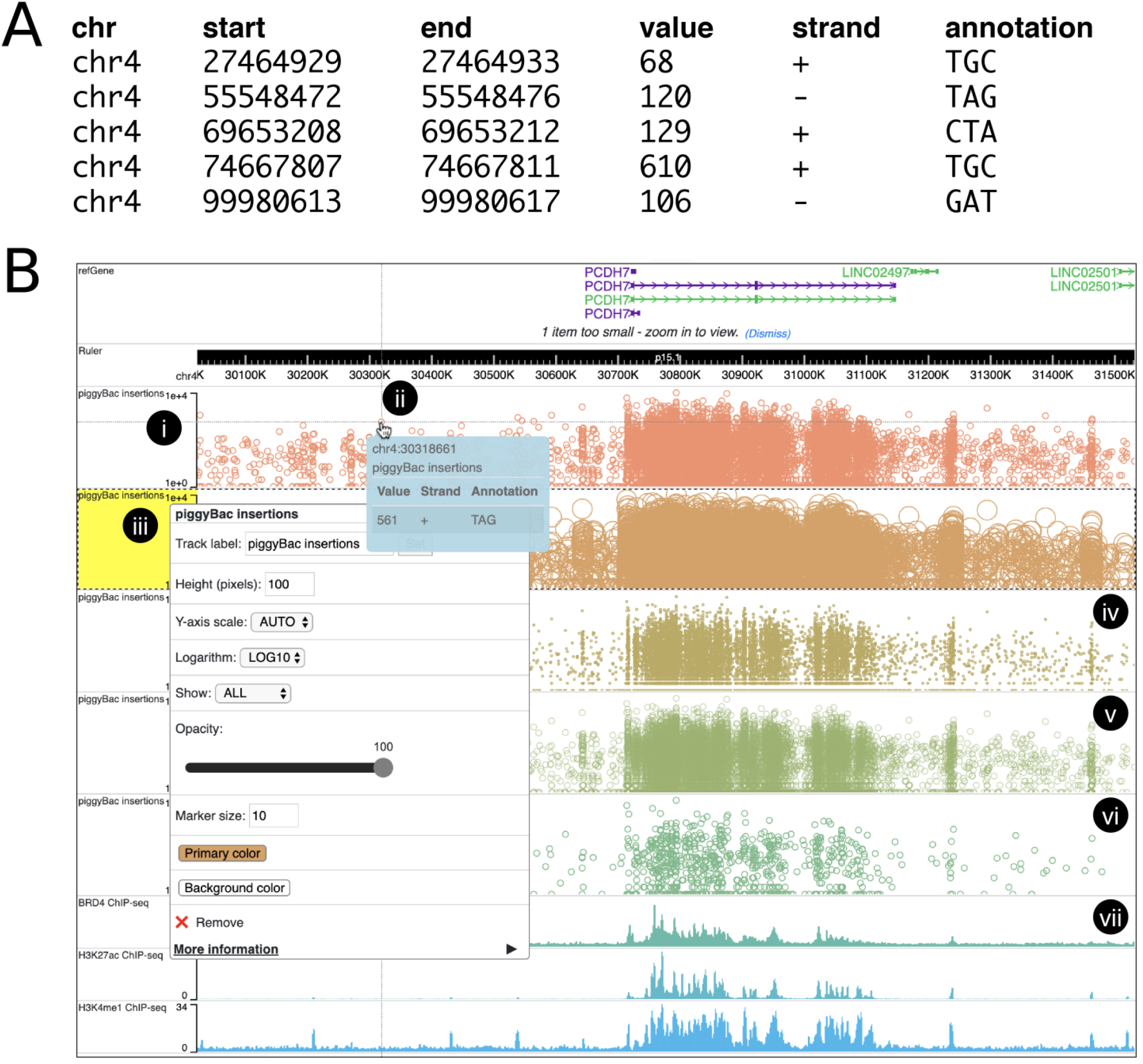
Overview of the qBED format and qBED tracks. (A) Example of a qBED file encoding transposon calling card data. The first three columns are inherited from the BED standard and encode the location of the insertion site. The fourth column stores the number of reads observed for each entry, while the fifth denotes strand. The sixth and final column is an annotation recording the sample-specific barcode for each insertion in the library. (B) Screenshot of qBED tracks depicting calling card data in the WashU Epigenome Browser. (i) qBED features appear on two-dimensional tracks, with genomic position along the *x*-axis and a numerical value on the *y*-axis (here, log-transformed read counts). (ii) An informational panel appears upon rollover of a calling card insertion, revealing read count, strand, barcode, and approximate location. (iii) Right-clicking on a qBED track pulls up a configuration panel. Tracks can be customized with respect to color, size, *y*-axis limits and transformations, marker size (iii-iv), opacity (v), and sample size (vi). (vii) Transcription factor and histone ChIP-seq data can be directly displayed alongside calling card data. [Link to this visualization.]

To visualize qBED files, we have created the qBED track and implemented it in the WashU Epigenome Browser (Li et al., 2019; Zhou et al., 2011), a leading portal for analyzing epigenomic data such as ChIP-seq, ATAC-seq, and Hi-C. (Prior to version 51.0.3, the qBED track was known as the calling card track). qBED tracks display circular markers for genomic features in two dimensions: genomic position along the *x*-axis and numerical value along the *y*-axis (Figure 1Bi). For calling card experiments, these represent transposon insertions and read counts, respectively. When the insertion coordinate spans more than one base, the marker is drawn at the midpoint of the interval. Moreover, as multiple insertions may occur at the same insertion site (e.g. from different replicates), multiple markers can co-occur at the same *x*-coordinate and stratify across the *y*-axis. qBED tracks support interactive exploration of data. As a cursor approaches a data point, a rollover pane appears (Figure 1Bii), displaying the read count, strand, and annotation (columns 4, 5 and 6, respectively). Near the top of the rollover pane is the track name and an approximate (to the nearest pixel) genomic location.

Right-clicking on a qBED track leads to a customization panel (Figure 1Biii). Individual tracks can be shaded in any RGB color (Figure 1Biii-vi), to better delineate different samples. The size of the calling card marker can be made larger (Figure 1Biii) or smaller (Figure 1Biv), depending on user preference. The opacity of the track can also be adjusted (Figure 1Bv), which may help reveal structure in regions of pronounced insertion density. For very large datasets, a random subsample of the data can be displayed (Figure 1Bvi). This prevents overplotting of markers and can reduce the browser’s memory consumption. Finally, and most importantly, the WashU Epigenome Browser enables calling card data to be natively visualized alongside other genomic datasets, such as ChIP-seq from the same cell type (Figure 1Bvii).

## Applications

qBED files present genomic data as a discrete point process as opposed to a pseudo-continuous function of sequencing coverage. In addition to analyzing calling card experiments, this format may also be useful for existing genomic data types. Here we present two such examples. Combined Annotation Dependent Depletion (CADD) scores integrate multiple streams of information to predict the deleteriousness of single nucleotide polymorphisms (SNPs) and indels (Kircher et al., 2014; Rentzsch et al., 2019). These are typically displayed as vertical lines depicting the mean and variance of scores observed for each base (Figure 2A). This approach, while useful as a summary statistic, does not allow for interactive exploration of individual mutations. We converted CADD scores for indels from variant call format (VCF) to a qBED file, using the numeric column to store the CADD score and the annotation column to store the mutation. When viewed on the WashU Epigenome Browser, individual polymorphisms can be inspected. A view of the homeobox gene *CRX* reveals a cluster of strongly deleterious indels in the terminal exon (Figure 2A). The qBED display emphasizes the density of variants along both the genomic (*x*) and CADD (*y*) axes, offering an unvarnished look at the complete spectrum of deleteriousness in a dataset.

**Figure 2:**
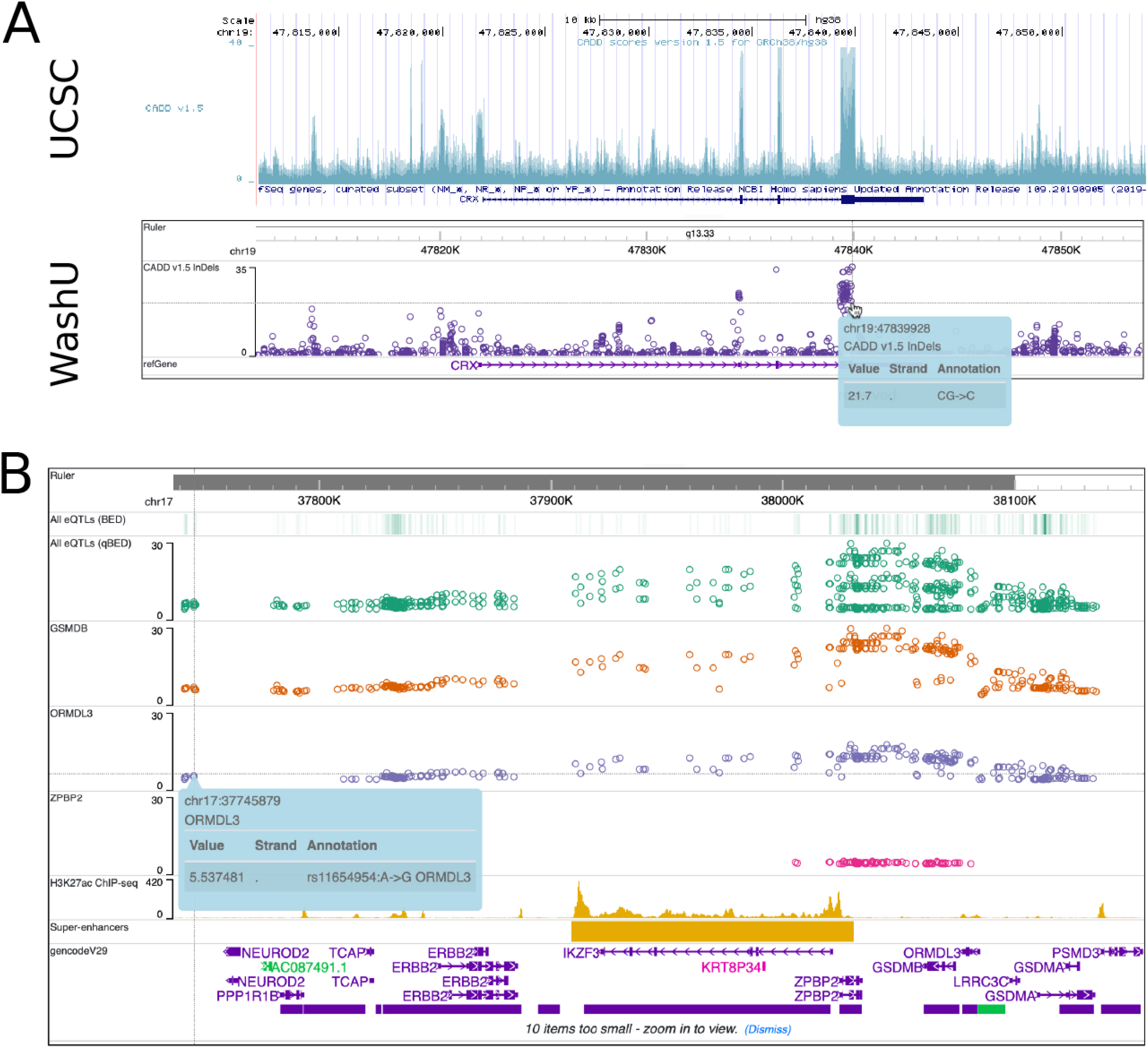
Application of the qBED specification to other genomic datasets. (A) Top: CADD scores for the gene *CRX,* as visualized on the UCSC Genome Browser. Bottom: CADD scores visualized on the WashU Epigenome Browser after conversion to qBED. Genomic position is along the *x*-axis and Phred-style CADD scores are along the *y*-axis. The mouseover pane reveals more information on an individual variant. [Link to this visualization.] (B) eQTLs for CD20+ B cells visualized as calling card tracks. The top track shows all significant eQTLs in view plotted as a BED (density) track, followed by a qBED representation of the same data. The *y*-axis represents the negative base-ten logarithm of the *p-*value. The next three tracks show significant eQTLs for the genes *GSDMB, ORMDL3,* and *ZPBP2,* respectively. Finally, we show H3K27ac ChIP-seq (coverage on the *y*-axis) and a super-enhancer for this cell type. A mouseover pane can reveal further details stored in the qBED file, including Reference SNP ID and mutation. [Link to this visualization.]

A second application of qBED files is in genome wide association studies (GWAS) and expression quantitative trait locus (eQTL) mapping, which aim to identify SNPs that are significantly correlated with either phenotypes or gene expression, respectively. Most significant SNPs fall in noncoding regions and their functional significance can be unclear (Gloss and Dinger, 2018; Tak and Farnham, 2015). One way to prioritize variants is by considering their regulatory and epigenetic context (Gloss and Dinger, 2018; Tak and Farnham, 2015); however, a quantitative view of SNPs is not supported by most genome browsers.

Investigators either manually align separate views of SNPs with views of epigenetic profiles, or encode SNPs as BED tracks, which shows position but sacrifices the quantitative measure-usually the negative base-ten logarithm of the *p*-value-of the association (Farh et al., 2015).

We reasoned that the qBED track could display both the density and the quantitative value of SNPs in association studies. We used a publicly available eQTL dataset from CD20+ B cells (Schmiedel et al., 2018) and converted it to qBED format, storing the negative base-ten logarithm of the *p*-value of the eQTL association in the numeric column; and storing the reference SNP, mutation, and linked gene in the annotation field. We simultaneously plotted H3K27ac ChIP-seq data (Davis et al., 2018; The ENCODE Project Consortium, 2012) and a track of super-enhancers for the same cell type (Figure 2B). Such data would either have to be manually aligned with another browser shot or plotted as a BED track (shown) that only emphasizes the local density of variants. The qBED visualization shows both the density of variants and the significance of each variant, alongside epigenetic context, all in a single pane. We can also separate eQTLs by target gene and assign them to individual tracks, revealing how genes in close proximity to each other can have different eQTL effect sizes from the same genomic sequence. In particular, eQTLs associated with *GSDMB* and *ORMDL3* expression span a large swath of flanking DNA, including overlapping an adjacent super-enhancer, while eQTLs associated with *ZPBP2* expression are constrained to a much narrower segment.

## Conclusion

The qBED specification and the accompanying qBED track offer researchers the ability to visualize genomic point processes-such as transposon insertions, polymorphism deleteriousness, or phenotypic associations-by adding a numerical *y-*axis for stratifying features on the genomic *x*-axis. We envision investigators using this format not only for analyzing calling card experiments, but any data involving relatively small, quantitatively separable genomic features. While we feel the six-column format presented here is complete enough for existing analyses, we leave open the possibility for future enhancements. In particular, extra columns could be added to encode secondary and/or tertiary information for each entry. These could be visualized, pending browser support, with either a numerical color scale, in the case of quantitative data, or different marker shapes, for categorical data.

## Acknowledgements

AM was supported by NIH F30 HG009986. RDM was supported by NIH RF1 MH117070 and R01 GM123203. T.W. was supported by NIH R01 HG007354, R01 HG007175, R01 ES024992, U01 CA200060, U24 ES026699, and U01 HG009391; and American Cancer Society RSG-14-049-01-DMC.

